# Measurement Equivalence of On-Scalp OPM-MEG and Cryogenic MEG for Auditory and Somatosensory Cortical Mapping Across Development

**DOI:** 10.64898/2026.06.01.728862

**Authors:** William Gaetz, George O’Neill, Charlotte Birnbaum, Teresa Cheung, Timothy P. L. Roberts, Jeramy Hughes, Svenja Knappe, Orang Alem

## Abstract

Wearable optically pumped magnetometer magnetoencephalography (OPM-MEG) reduces sensor-to-cortex distance compared with conventional cryogenic SQUID-MEG, but whether the two technologies yield equivalent neurophysiological conclusions remains unclear. We recorded auditory and somatosensory evoked fields in 18 participants (10–45 years) using a 128-sensor FieldLine HEDscan OPM-MEG system and a 275-channel CTF SQUID-MEG system within the same individuals. Equivalent current dipole source models were estimated using identical preprocessing and modeling procedures and compared using paired permutation testing.

Both systems localized canonical auditory and somatosensory cortical generators with matched peak latencies and modest cross-system spatial differences. Auditory sources showed a consistent medial bias in SQUID-MEG localization, whereas somatosensory sources exhibited a small systematic offset (∼4 mm), indicating stable coordinate differences rather than localization error. Dipole moments were larger for SQUID-MEG and goodness-of-fit higher for OPM-MEG; however, the increased moment was explained by a medial localization bias, demonstrating inverse-model effects rather than physiological disagreement. Auditory dipole moment increased with age in both systems, whereas somatosensory responses showed no age-related change. Together, these observations indicate preserved developmental physiology across platforms.

These findings demonstrate that OPM-MEG and SQUID-MEG recover the same cortical generators and support equivalent biological interpretations despite predictable geometry-dependent coordinate differences. OPM-MEG therefore represents a measurement-equivalent implementation of MEG suitable for sensory functional mapping.

## Introduction

Magnetoencephalography (MEG) measures neuronal currents with millisecond precision, but it remains unclear whether systems based on fundamentally different sensor physics recover equivalent cortical generators or instead produce platform-dependent physiological conclusions. Conventional cryogenic systems based on superconducting quantum interference devices (SQUIDs) require fixed helmets and liquid helium cooling, which constrain motion and limit applicability in children [1, 2]. Optically pumped magnetometers (OPMs) enable wearable, room-temperature MEG with sensors positioned directly on the scalp [3–5]. Early OPM-MEG studies demonstrated the feasibility of recording canonical evoked responses using room-temperature sensors [6, 7]. A small number of direct OPM–SQUID comparison studies have reported broadly similar evoked-field patterns, but systematic within-subject comparisons remain sparse, particularly across a broad developmental age range [8, 9]. As a result, it remains uncertain whether differences in sensor placement, coverage, and spatial sampling could introduce systematic differences in estimated source localization or amplitude.

To address this question, we recorded auditory and somatosensory evoked responses from 18 participants (children and adults) using both a 128-sensor FieldLine Medical HEDscan OPM system and a 275-channel CTF cryogenic gradiometer system in a within-participant design. We quantified cross-platform agreement in latency, amplitude scaling, and source localization, and examined age-related effects within and across platforms. Because sensor-to-cortex distance varies with head size in fixed-helmet MEG, we asked whether age-dependent scaling effects would be reduced in OPM recordings. The objective of this study was therefore to determine whether OPM-MEG and SQUID-MEG recover the same cortical generators and physiological interpretations despite differences in sensor geometry and field sampling.

## Methods

### Subjects

Eighteen participants (14 female; age range 10.9–45.5 years, mean ± SD = 23.9 ± 11.1 years) were included in the final analysis. All procedures were approved by the Children’s Hospital of Philadelphia (CHOP) institutional IRB. Written informed consent was obtained from adult participants and from parents or guardians of minors. Assent was obtained from all participating children.

### Data acquisition

All participants completed both experiments across two sessions on the same day. Data were recorded first on a conventional cryogenic CTF MEG system (275-channel gradiometer sensors; CTF, Port Coquitlam, BC) and then using the HEDscan OPM system (128 single-axis magnetometer sensors; FieldLine Medical, Louisville, CO). For the OPM recordings, sensors were mounted in the HEDscan Smart Helmet™ (FieldLine Medical, Louisville, CO) that positions the sensors directly on the scalp and conforms the array to each participant’s head size and shape. The Smart Helmet was then secured to a mounting structure to maintain a seated position comparable to that used during CTF-MEG acquisition. The sensor positions and orientations are determined automatically by the HEDscan system for each subject, and coregistered to anatomy using head-position measurements.

### Common anatomical reference frame

Three fiducial locations (nasion, left preauricular, right preauricular) were first marked directly on the participant’s skin. Head-position indicator (HPI) coils were then affixed over these same marked locations and remained in place for both recordings, ensuring identical physical fiducial positions across the CTF and HEDscan sessions. Each MEG system tracked the coils using its native signal generation software method, but critically, the underlying coil positions were unchanged between systems. The same anatomical landmarks were subsequently overlaid with MRI-visible markers to allow alignment to each participant’s structural MRI. This procedure established a common anatomical reference frame for both MEG datasets rather than independent co-registrations.

### Experimental Stimuli Auditory

Auditory stimuli consisted of 1000-Hz pure tones (300 trials) presented with a variable inter-stimulus interval (1 s ± 500 ms) via a centrally positioned piezoelectric speaker. Preprocessing and averaging followed the same pipeline described above.

### Somatosensory

Pneumatic pulses were delivered separately to left and right index fingers using clip-on balloon diaphragms (30 psi, 35 ms duration, ISI = 0.75–0.95 s). Each participant completed 500 trials per hand. Preprocessing and averaging followed the same procedure used for the auditory data.

### MEG Data Preprocessing

Each system used its native noise-suppression strategy. The CTF data employed reference sensors and third-order synthetic gradiometry [10] to remove external interference. The HEDscan data were denoised using Adaptive Multipole Modelling (AMM) [11], which separates fields originating inside versus outside the sensor array, projecting external components out of the data.

Following denoising, both datasets underwent identical preprocessing: removal of persistently noisy channels, notch filtering at 60 Hz and 25 Hz (to address the idiosyncratic environmental noise), band-pass filtering from 3–40 Hz, epoching around stimulus onset (-250 to 300 ms), and baseline correction using the pre-stimulus interval. Noise covariance matrices were computed from baseline data (-250 to 0 ms) across all trials within each participant and used for signal-to-noise ratio estimation and source modeling.

### Equivalent Current Dipole (ECD) Modelling

Source localization was performed using maximum-likelihood ECD fitting [12] in FieldTrip [13]. Averaged data were whitened using the corresponding noise pre-stimulus data covariance [14]. For auditory stimuli, a pair of symmetric bilateral dipoles were fitted to a 10 ms post-stimulus window, capturing the auditory M100 response. The window centering was subject-specific (range: 90-135 ms after stimulus onset). The dipole fit consisted of two stages; a coarse grid search (15 mm spacing between locations) identified the best fit regions, followed by a gradient descent optimization of the final dipole location parameters. A symmetry constraint along the longitudinal fissure was applied during both the initial search and final optimization. Dipoles were modeled using Nolte’s single shell [15], where the conductive model geometry was based on a convex hull mesh of the brain/CSF boundary, derived from the individual subject’s MRIs.

For somatosensory stimuli, a single stationary dipole was fit within the 80–120 ms post-stimulus window, corresponding to the contralateral P50m response in primary somatosensory cortex (SI). Again, a coarse grid search followed by a fine gradient descent optimization was applied to determine the final dipole location parameters.

### Statistical Testing

To determine whether the two recording systems produced comparable results, we used a paired permutation framework appropriate for matched measurements. For each observation, a difference score was computed between systems (e.g., goodness-of-fit, reconstructed time-course metrics), preserving the within-subject and within-event correspondence. Under the null hypothesis of no systematic difference between systems, the sign of each paired difference is arbitrary.

For source localization coordinates, paired differences were treated as 3-dimensional displacement vectors between systems. Under the null hypothesis of no systematic localization bias, the direction of each paired displacement vector is arbitrary. We therefore generated a null distribution by randomly flipping the sign of each displacement vector across 5,000 permutations and recalculating the magnitude of the mean displacement vector at each iteration. The observed mean displacement magnitude was compared with this distribution to obtain a two-tailed p-value.

This approach treats each homologous source pair (subject × hemisphere) as the exchangeable unit and avoids assumptions of independence between hemispheres or repeated measurements within a subject. For scalar paired measures (e.g., latency, goodness-of-fit, and dipole moment), paired sign-flip permutation tests were applied to the mean difference between systems.

## Results

### Auditory Evoked Field (AEF) Localization

Group-averaged dipole localizations for the auditory evoked fields are shown in Figure 3. All sources localized to the superior temporal plane and were oriented approximately perpendicular to the cortical surface, consistent with activation of primary auditory cortex.

**Figure 1.**
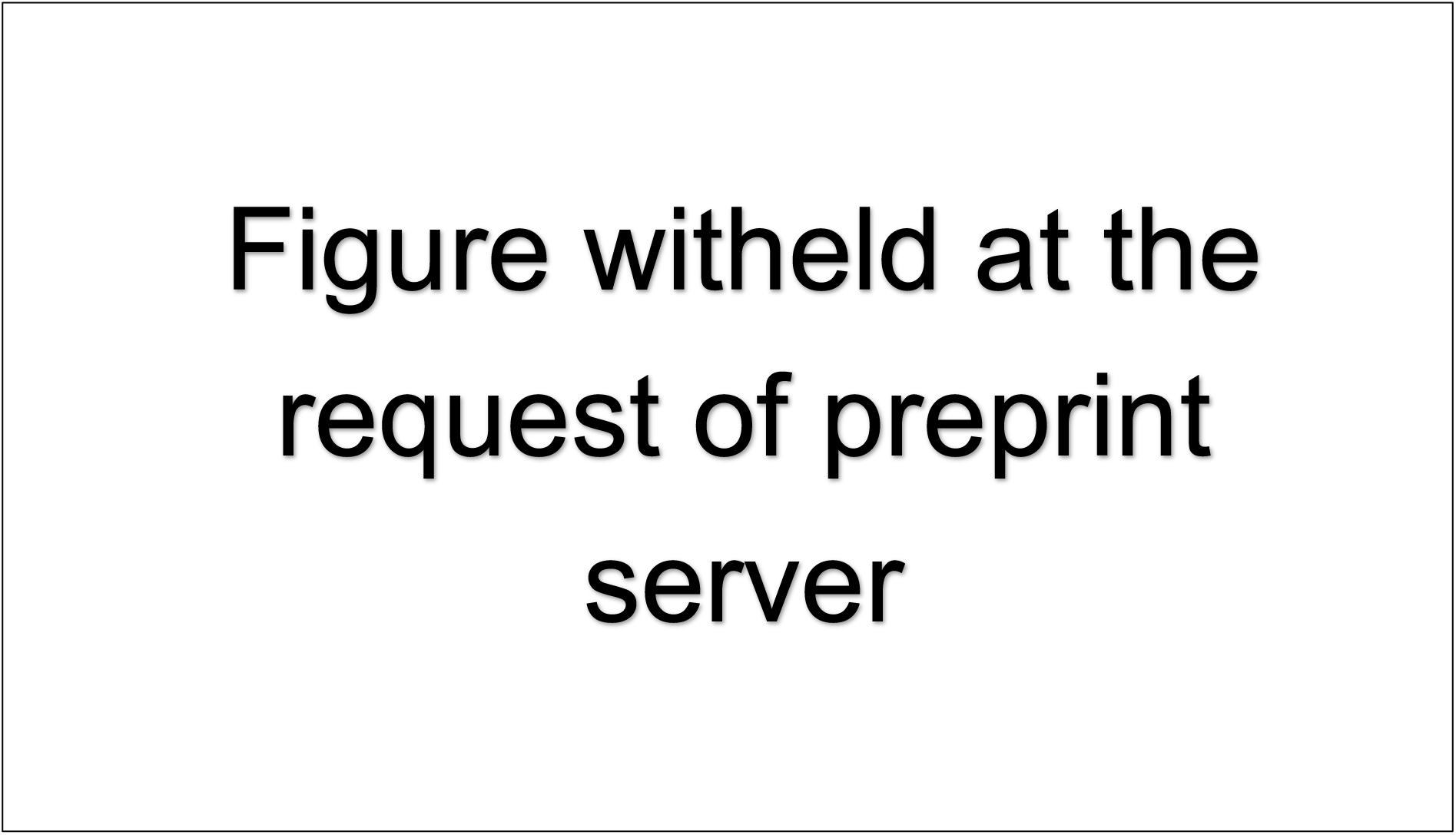
HEDscan 128-channel OPM system (left) and CTF-Omega 275 channel systems (right).

**Figure 2.**
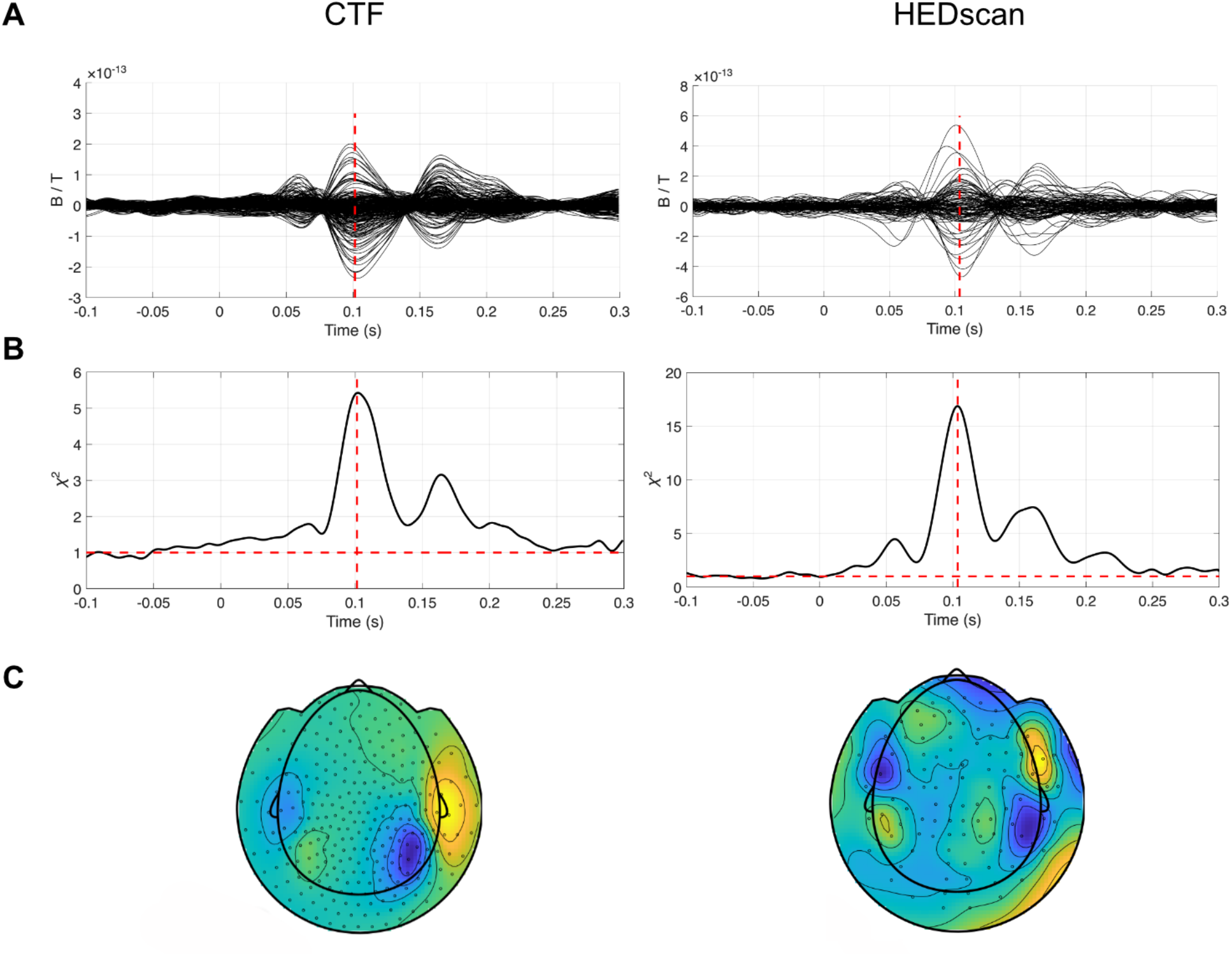
Representative Auditory Evoked Response and Topoplot from CTF (left column) and HEDscan (right) from a single participant. **A)** Butterfly plots of the trial-averaged evoked response. Dashed vertical lines represent the timepoint used to fit the dipoles for each system (101.4 ms for CTF v 103.6 ms for HEDscan). **B)** Global Field Power (GFP) of data pre-whitened against the pre-stimulus covariance, red horizontal dashed line represents a signal-to-noise ratio of 1. The correlation between the two system’s GFP time series is 0.979. **C)** Topoplots of the evoked response at the selected latencies (101 ms and 103 ms respectively), showing both systems capture asymmetry of the strength of the evoked responses in this participant’s auditory cortices.

**Figure 3.**
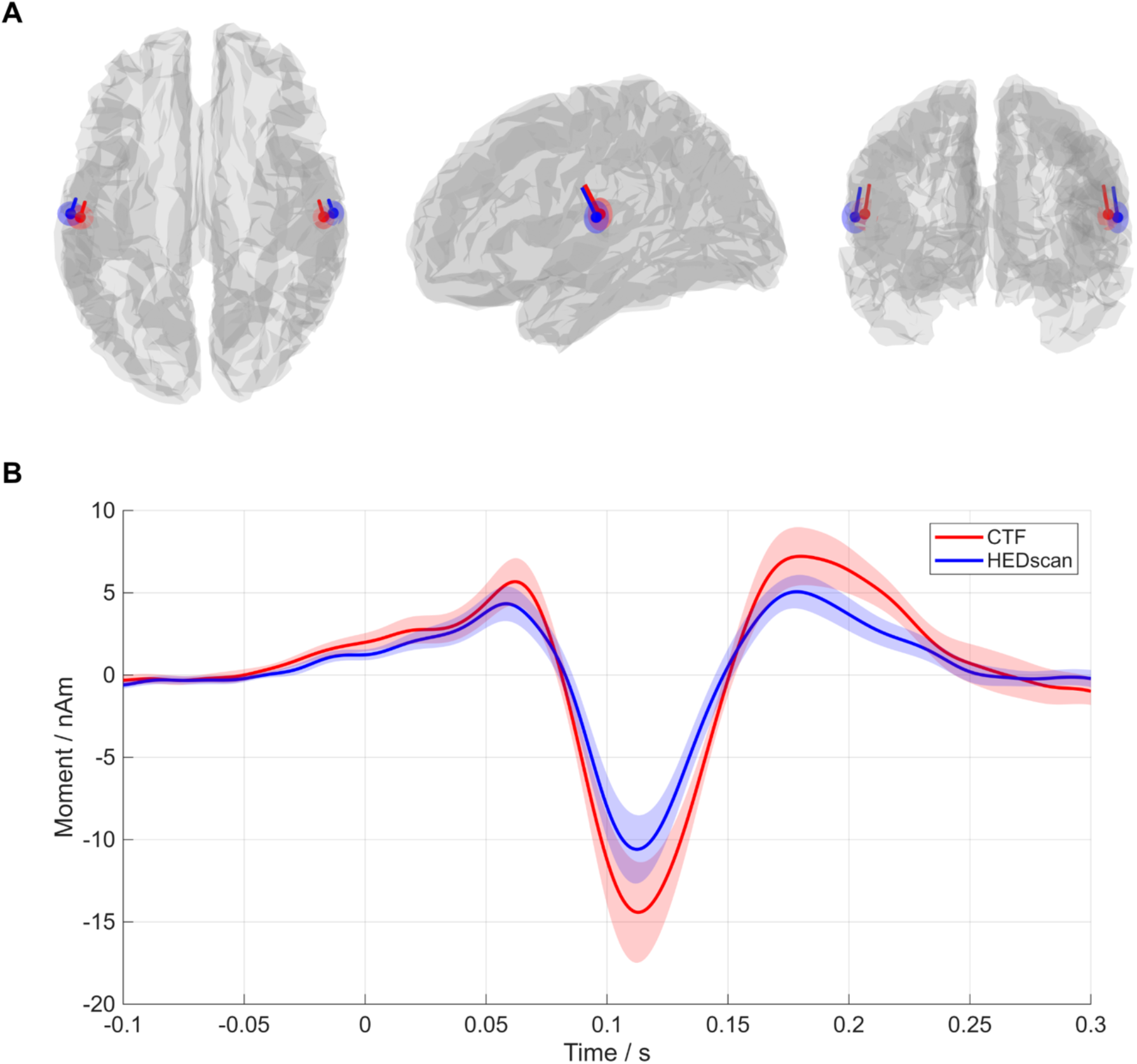
Group results from the equivalent current dipole fitting on the binaural auditory stimulation. **A)** Average fitted dipole locations for left and right auditory sources for the CTF MEG system (red) and HEDscan system (blue) represented on a cortical sheet. Solid points represent the location of the dipole, lines represent the orientation, shaded areas represent 95% confidence boundaries for the localization. **B)** The group-averaged time-courses of the auditory evoked signals observed from the ECD source locations. Solid lines represent the average of all sources for a given MEG system, and the shaded areas are the standard errors over the mean. Permutation testing across whole time series reveals no significant effects between systems at any timepoint. Group-averaged dipole localizations for the somatosensory-evoked fields are shown in Figure 5. All sources localized to the central sulcus and were oriented perpendicular to the cortical surface, consistent with activation of the primary somatosensory cortex. Detailed source coordinates in Montreal Neurological Institute (MNI) space are provided in Table 2.

**Figure 4.**
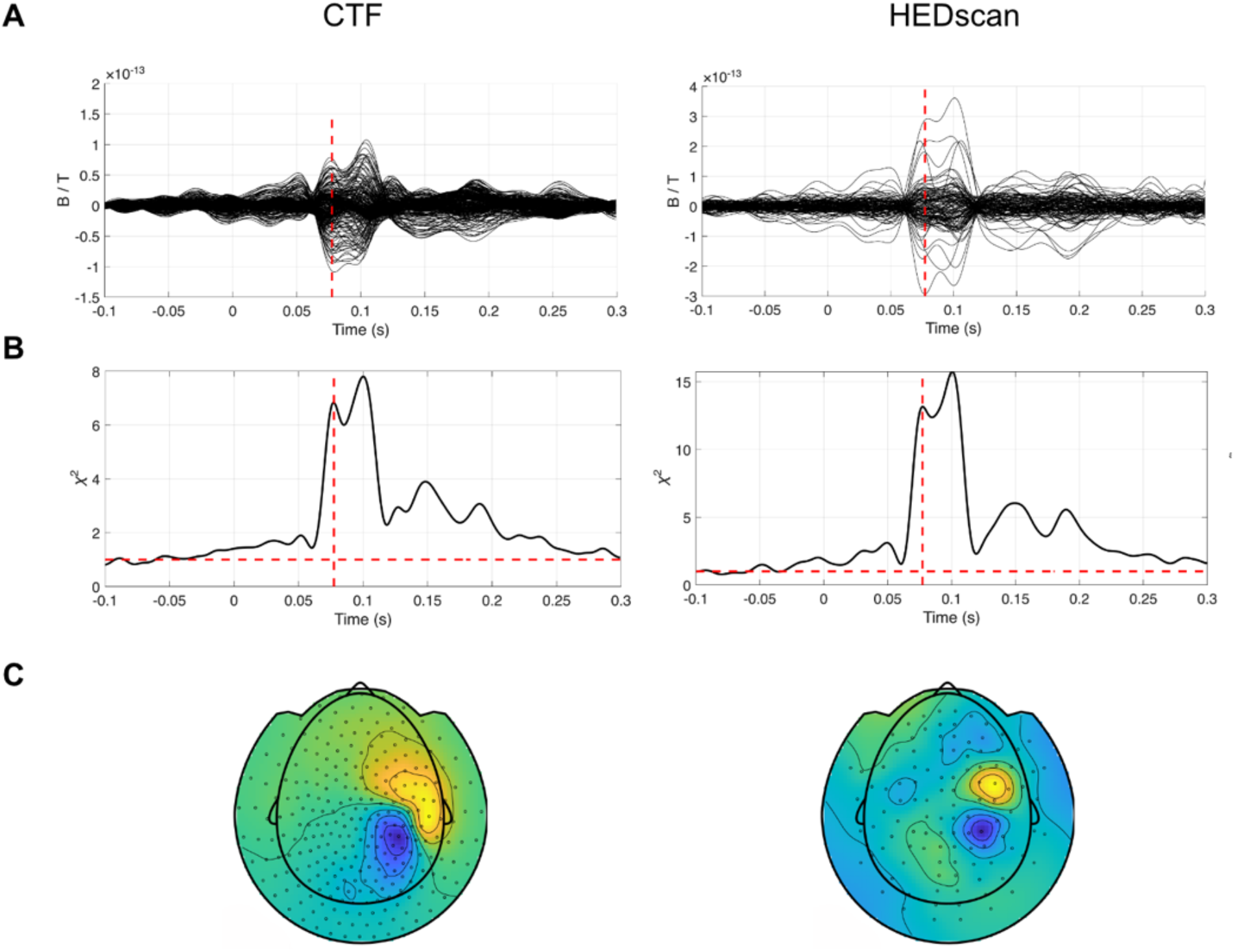
Representative Somatosensory-Evoked Response and Topoplot from CTF (left column) and HEDscan (right) from a single participant. **A)** Butterfly plots of the trial-averaged evoked response. Dashed vertical lines represent the timepoint used to fit the dipoles for each system (77.4 ms for CTF v 77.3 ms for HEDscan). **B)** Global Field Power (GFP) of data pre-whitened against the pre-stimulus covariance, red horizontal dashed line represents a signal-to-noise ratio of 1. The correlation between the two systems’ GFP time series is 0.993. **C)** Topoplots of the evoked response at the selected latencies (77 ms).

### Overall localization agreement for AEF dipoles

We first asked whether the two systems localized the *same cortical point* on average, or whether one system consistently placed sources in a different region of the brain. To test this, for each homologous source pair (subject × hemisphere) we computed the signed 3-dimensional displacement vector between systems and evaluated whether the mean displacement differed from zero using a paired sign-flip permutation test (5,000 permutations). No significant overall offset was observed (magnitude of the mean signed displacement vector 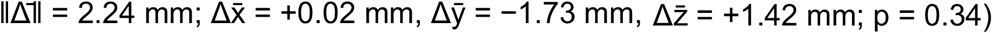 indicating that neither system systematically localized auditory cortex to a different anatomical location. Because mediolateral shifts reverse sign across hemispheres, averaging signed displacement vectors across hemispheres can reduce the magnitude of the mean vector even when consistent hemisphere-specific offsets are present.

### Cross-system AEF source separation

We next quantified the magnitude of the spatial difference between paired CTF and HEDscan auditory dipole estimates, independent of direction. For each homologous source pair, we computed the absolute Euclidean distance between the two system estimates. The absolute separation between system estimates averaged 11.97 mm (median 11.37 mm; range 3.61–22.88 mm), which is within the same general scale as previously reported test–retest variability and intrasubject dispersion of MEG equivalent current dipole localization, rather than indicating gross cross-platform disagreement [16-18].

### Directional bias (axis-specific behavior)

Although no overall bias was present, we next asked whether the displacement followed a consistent anatomical direction that could cancel out in the average. Hemisphere-specific analysis revealed a small medial shift of the CTF dipoles relative to HEDscan in both hemispheres (Left: 4.79 mm, p = 0.048; Right: 4.76 mm, p = 0.058). Thus, while overall localization agreement was preserved, the cryogenic system tended to place the auditory source slightly closer to the midline.

To determine whether the hemisphere-dependent displacement reflected a consistent directional difference rather than random variation, the mediolateral component was tested after correcting for hemispheric sign reversal (left and right hemispheres shift in opposite coordinate directions for the same medial movement). A paired sign-flip permutation test demonstrated a systematic medial shift of CTF dipoles relative to HEDscan dipoles (mean medialization = 4.22 mm, p = 0.00020). Thus, despite a small mean displacement across paired observations, AEF localization demonstrated a consistent cross-system mediolateral difference across hemispheres, with CTF dipoles localized more medially than HEDscan dipoles. The association between mediolateral displacement and dipole moment suggests that the observed amplitude differences are at least partly attributable to localization geometry rather than physiological variation. Thus, although the two systems localized the same auditory generator, the relative medial displacement of the cryogenic-system estimates was associated with predictable scaling of estimated source strength.

### AEF ‘N100m’ Dipole Latency, Fit Quality (GOF), and Strength (Moment)

No detectable differences in AEF peak latency were observed between systems (mean difference = +1.86 ms, p = 0.219). Goodness-of-fit values did not differ between systems (mean difference CTF – HEDscan = −0.025, p = 0.244). In contrast, dipole moments were systematically larger for CTF (mean log|moment| difference = +0.380, p = 0.00080). The difference in dipole moment between systems was strongly associated with the mediolateral displacement (Spearman ρ = 0.63, p = 0.0006), indicating that larger CTF source amplitudes occurred in subjects where the dipole was localized more medially.

### Age-related effects

Auditory dipole moment increased with age in both systems (CTF: ρ = 0.38, p = 0.026; HEDscan: ρ = 0.47, p = 0.0032). A paired sign-flip permutation test on the within-subject slope differences revealed no difference in the age–moment relationship between systems (p = 0.66).

### Somatosensory-Evoked Field (SEF) Localization

Group-averaged dipole localizations for the somatosensory-evoked fields are shown in Figure 5. All sources localized to the central sulcus and were oriented perpendicular to the cortical surface, consistent with activation of the primary somatosensory cortex. Detailed source coordinates in Montreal Neurological Institute (MNI) space are provided in Table 2.

**Table 1.** Hemisphere-specific paired localization offsets between systems (AEF). Mean displacement vectors (CTF − HEDscan) computed from paired subject-level dipole localizations within each hemisphere. P-values derived from paired sign-flip permutation testing (5,000 permutations).

| Hemisphere | Mean $\Delta x$ (mm) | Mean $\Delta y$ (mm) | Mean $\Delta z$ (mm) | Offset magnitude (mm) | p-value |
| --- | --- | --- | --- | --- | --- |
| Left AEF | +4.25 | -1.71 | +1.41 | 4.79 | 0.048 |
| Right AEF | -4.20 | -1.74 | +1.43 | 4.76 | 0.058 |

**Table 2.**
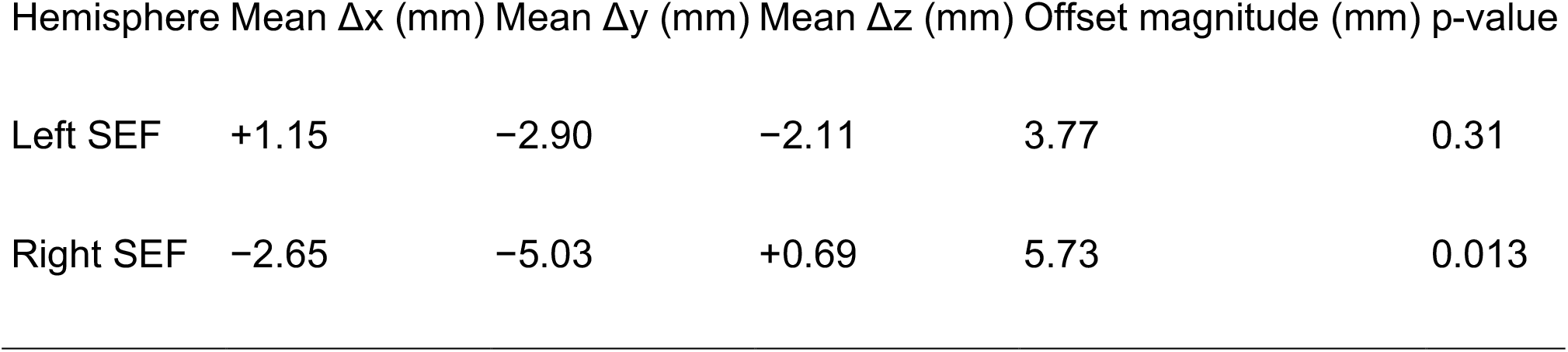
Hemisphere-specific paired localization offsets between systems. Mean displacement vectors (CTF − HEDscan) computed from paired subject-level dipole localizations within each hemisphere. P-values derived from paired sign-flip permutation testing (5,000 permutations).

**Figure 5.**
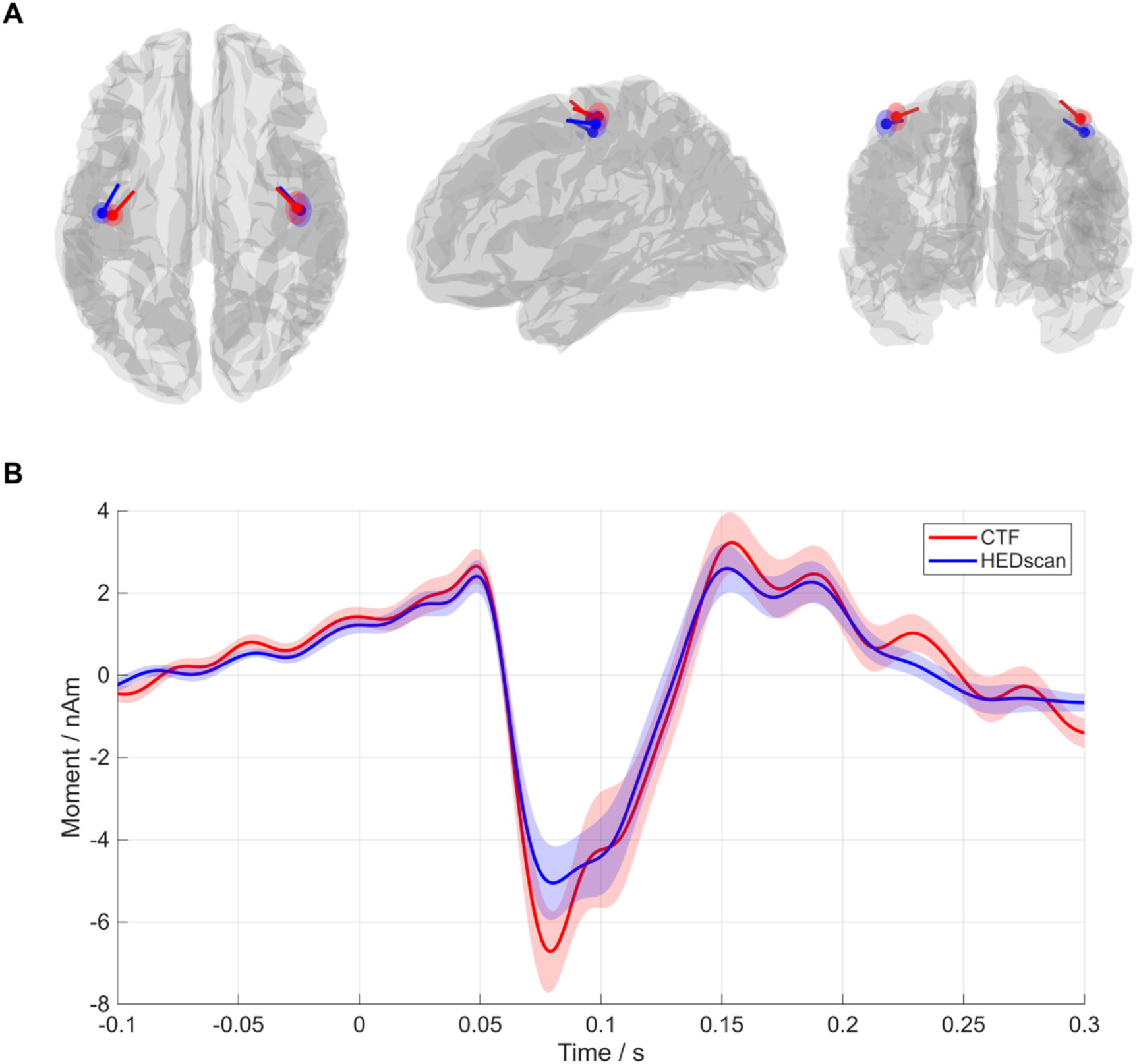
Group results from the equivalent current dipole fitting on the somatosensory stimulation data. **A)** Average fitted dipole locations for left and right somatosensory sources for the CTF system (red) and HEDscan system (blue) represented on a cortical sheet. Solid points represent the location of the dipole, lines represent the orientation, shaded areas represent confidence boundaries for the localization. **B)** The group-averaged time-courses of the somatosensory-evoked responses, derived from the ECD source locations. Solid lines represent the average of all sources for a given MEG system and the shaded areas are the standard errors over the mean. Permutation testing across whole time series reveals no significant effects between systems at any timepoint.

### Overall localization agreement for SEF dipoles

We first evaluated whether the two recording systems produced systematic differences in source localization. For each homologous source pair (subject × hemisphere), the signed three-dimensional displacement vector between systems was computed and tested against zero using a paired sign-flip permutation test (5,000 permutations). A small but significant mean offset was observed (mean displacement magnitude = 4.10 mm; Δx = −0.75 mm, Δy = −3.97 mm, Δz = −0.71 mm; p = 0.028).

When examined separately, the spatial offset was significant in the right hemisphere (mean offset 5.73 mm, p = 0.013) but not in the left hemisphere (mean offset 3.77 mm, p = 0.31). For reference, the absolute Euclidean distance between paired dipole localizations averaged 12.00 mm (median 10.20 mm; range 2.46–30.54 mm). Of note, this measure reflects both measurement variability within each system and any true spatial offset between systems.

### Cross-system SEF source separation

We next quantified the magnitude of the spatial difference between paired CTF and HEDscan somatosensory dipole estimates, independent of direction. For each homologous source pair, we computed the absolute Euclidean distance between the two system estimates. The absolute separation between system estimates averaged 12.00 mm (median 10.20 mm; range 2.46–30.54 mm). This value reflects the overall spatial difference between paired system estimates and does not distinguish whether the separation arises from measurement variability within either system or from a stable offset between systems.

### Directional bias analysis

To determine whether the somatosensory displacement reflected a consistent anatomical direction, axis-specific paired permutation tests were performed. The displacement showed a significant posterior shift of CTF sources relative to HEDscan (mean Δy = −3.97 mm, p = 0.0026), significant in the right hemisphere (Δy = −5.03 mm, p = 0.0036) but not in the left (p = 0.153). No consistent mediolateral bias was observed.

### SEF ‘P50m’ Dipole Latency, Fit Quality (GOF), and Strength (Moment)

No detectable temporal offset between systems in SEF peak latency was observed (mean difference < 0.001 ms, p = 0.98). We next compared dipole fit characteristics between systems using paired permutation testing. Goodness-of-fit values were significantly higher for HEDscan than CTF (mean difference CTF − HEDscan = −0.092, p = 0.0002), indicating better model fit values for HEDscan. In contrast, dipole moments were systematically larger for CTF (mean logarithmic absolute difference = +0.208, p = 0.008). Unlike the auditory responses, SEF dipole moment did not correlate with displacement magnitude, indicating that the offset does not influence forward-model scaling.

### Age-related effects

No significant associations were observed between age and SEF dipole latency, goodness-of-fit, or dipole moment in either recording system (all p > 0.12). A weak, non-significant trend toward increasing goodness-of-fit with age was observed in HEDscan (ρ = 0.32, p = 0.054).

The spatial offset did not influence dipole moment or latency, indicating that the displacement reflects a stable coordinate frame difference rather than a change in the estimated cortical generator. In contrast to the auditory responses, the somatosensory data supported measurement equivalence across platforms: both systems recovered the same physiological source despite a small systematic spatial translation.

## Discussion

This study evaluated whether OPM-MEG and SQUID-MEG recover the same cortical generators and physiological interpretations despite differences in sensor geometry and field sampling. Both platforms reproduced canonical auditory and somatosensory generators with closely matched peak latencies and similar dipole locations. Across most physiological measures, the systems were concordant, although small modality-specific differences in localization and inverse-model geometry were observed. Overall, the results indicate that the two systems measure the same underlying neural activity despite differences in sensor technology. The magnitude of spatial offsets was within the same general scale as typical test–retest dipole variability reported for clinical MEG. Operationally, measurement equivalence in this context refers to preservation of source location, temporal characteristics, and developmental associations across platforms, even when absolute amplitude scaling or coordinate frames differ.

Localization agreement was high across both modalities. The auditory responses showed no consistent global displacement but exhibited a directional medial bias in the cryogenic system. The magnitude of cross-system spatial separation was modest, indicating a stable coordinate difference rather than gross disagreement in the underlying cortical generator. These findings suggest minor platform-specific coordinate and scaling differences rather than differences in the underlying cortical generators.

The larger dipole moments estimated in the CTF recordings were not independent of localization geometry. Consistent with the directional permutation analysis, AEF dipoles localized more medially in the CTF data, and the magnitude of this medial displacement strongly predicted the increase in estimated dipole moment. Previous work has shown that sensor coverage and source geometry can influence dipole localization and moment estimates in conventional MEG systems [19, 20], which is consistent with the possibility that such factors contributed to the directional effect observed here.

This relationship indicates that the amplitude difference primarily reflects geometric effects of source placement within the forward model rather than differences in neural activity or sensor sensitivity. In this dataset, a shift toward the midline was associated with increased lead-field gain, inflating the fitted moment required to explain the measured field. Thus, the amplitude discrepancy between systems arises from systematic spatial bias rather than platform-specific measurement scaling. In contrast, the somatosensory responses showed a different behavior. The SEF offset occurred primarily along the anterior–posterior axis and did not predict dipole moment differences. This dissociation indicates that the auditory shift reflects inverse modeling of bilateral generators collapsing toward the midline, whereas the somatosensory shift reflects a stable coordinate difference related to spatial sampling geometry rather than changes in estimated neural activity, consistent with differences in sensor spatial sampling between on-scalp OPM and fixed-helmet SQUID arrays. The differing behavior across modalities is consistent with complementary hardware properties. The OPM array benefits from proximity to cortex and improved field pattern fidelity, whereas the denser SQUID array (275 cryogenic gradiometers vs 128 single-axis OPM magnetometers) provides finer spatial sampling of the magnetic field. The somatosensory offset therefore likely reflects limited spatial resolution rather than generator mislocalization. The modest pairwise separation and consistent directional structure further indicate stable estimation of the same cortical generator across platforms. In practical terms, the two systems localize the same source within slightly shifted coordinate frames.

### Developmental Implications

Beyond overall cross-platform concordance, the age-related findings support preservation of the same developmental physiology across systems. Auditory dipole moment increased with age in both systems with comparable slopes, indicating that the developmental effect reflects underlying physiology rather than measurement platform. Prior EEG and MEG studies have similarly reported increasing auditory response amplitude across development [21, 22]. In contrast, somatosensory responses showed only a nonsignificant trend towards increasing goodness-of-fit with age. Because the moment difference was geometrically mediated, the age effect represents physiological maturation rather than platform sensitivity.

The medial shift observed in the cryogenic system may reflect spatial smoothing of bilateral auditory generators rather than improved sensitivity to deep sources. Bilateral synchronous activation in auditory cortex may be represented by a deeper or more medial equivalent dipole when spatial gradients are insufficiently resolved. By preserving lateral field gradients more faithfully, on-scalp OPM measurements may better constrain the inverse solution and reduce collapse toward a single medial dipole [23-25].

These developmental patterns suggest that auditory and somatosensory systems mature along distinct trajectories and that the differing developmental patterns across modalities reflect maturation of auditory versus somatosensory systems rather than measurement platform. Age-related differences were smallest in adults and more variable in younger participants. Because OPM arrays can conform to head shape, they may reduce age-dependent geometric biases inherent to fixed-helmet MEG designs. Future cross-platform work should incorporate head circumference or cortical surface distance as covariates, as these geometric factors directly influence signal-to-noise and localization accuracy.

From a methodological standpoint, the systems differ in sensor type and interference control. The HEDscan array measures magnetic fields as magnetometers with adaptive multipole modeling for post-hoc noise suppression, whereas the CTF system employs third-order synthetic gradiometry. Although magnetometers and gradiometers differ in their depth sensitivity and coupling to distant interference, both systems produced equivalent source locations and latencies once interference was removed and data were whitened. This convergence demonstrates that ECD modeling provides consistent localization across sensor technologies, although absolute amplitudes must be interpreted in light of differing transfer functions.

### Clinical and translational implications

The OPM platform reproduced time-locked sensory responses widely used in presurgical and developmental mapping. Its ability to record in children with improved comfort and reduced sensor distance supports its use as a practical clinical alternative to conventional MEG. Enhanced motion tolerance may also reduce the need for general anesthesia in young pediatric epilepsy cases. These findings support the promise of OPM-based MEG for pediatric clinical use and motivate further validation in naturalistic and functional mapping paradigms.

Several limitations should be noted. The OPM sensors were mounted on a rigid frame rather than a fully wearable cap, so these results likely reflect optimal OPM performance under motion-free conditions. Our sample included both children and adults but may be underpowered to reveal systematic differences in sensitivity in very young participants recorded with conventional adult cryogenic MEG hardware because of the larger sensor-to-cortex distance. We focused on ECD modeling rather than beamforming or distributed inverses and examined only early sensory paradigms.

Taken together, the results show that agreement between systems is governed primarily by neurophysiology rather than sensor technology. Differences between systems followed predictable physical constraints: coordinate offsets reflected geometry and spatial sampling, goodness-of-fit reflected sensor proximity, and dipole moment reflected lead-field scaling. However, the biological inferences were unchanged. Both systems localized the same cortical generators, preserved modality-specific behavior, and reproduced expected developmental patterns. Thus, rather than demonstrating simple similarity, these findings show convergent hardware implementations recovering the same neural sources within expected inverse-model uncertainty. In this sense, within the limits of ECD modeling, the remaining differences are best understood as predictable coordinate and scaling differences between systems, not disagreement about the underlying neural generators. Future work should evaluate fully wearable, child-sized caps; expand to dual- or triaxial sensors; and extend cross-platform comparisons to beamformer and distributed source reconstructions.

## Conclusions

OPM-MEG and SQUID-MEG recover the same cortical generators and support the same physiological interpretations despite predictable differences in sensor geometry and field sampling. Remaining differences were small and consistent with predictable effects of sensor geometry and inverse model geometry rather than differences in the underlying cortical generators.

These findings demonstrate cross-technology measurement equivalence: independent MEG hardware implementations recover the same neurophysiological signals. Accordingly, on-scalp OPM-MEG represents a measurement-equivalent implementation of MEG rather than a separate modality, providing a practical platform for sensory cortical mapping in populations where wearable recordings are advantageous.

## Acknowledgements

The authors thank all the volunteers who participated in this study. The authors acknowledge John Dell, Rachel Golembski, Erin Verzella, Peter Lam, Shivani Patel, and Na’Keisha Robinson (RTs) for technical assistance during the study visits. This study was supported by the National Institutes of Health SBIR Phase II project (R43MH118154) Center PI: O. Alem; CHOP PI: W. Gaetz. The work was also supported in part by NIH-R61MH135114.

## References

1. Hämäläinen MS (1992) Magnetoencephalography: a tool for functional brain imaging. Brain Topogr 5:95–102.

2. Schwartz ES et al. (2010) Magnetoencephalography. Pediatr Radiol 40:50–58.

3. Sander TH, Preusser J, Mhaskar R, Kitching J, Trahms L, Knappe S (2012) Magnetoencephalography with a chip-scale atomic magnetometer. Biomed Opt Express 3:981–990.

4. Borna A, Carter TR, Colombo AP, Jau YY, McKay J, Weisend M, Taulu S, Stephen JM, Schwindt PDD (2020) Non-invasive functional-brain-imaging with an OPM-based magnetoencephalography system. PLoS One 15:e0227684.

5. Alem O, Hughes KJ, Buard I, Cheung TP, Maydew T, Griesshammer A, Holloway K, Park A, Lechuga V, Coolidge C, Gerginov M, Quigg E, Seames A, Kronberg E, Teale P, Knappe S (2023) An integrated full-head OPM-MEG system based on 128 zero-field sensors. Front Neurosci 17:1190310.

6. Boto E, Meyer SS, Shah V, Alem O, Knappe S, Kruger P, Fromhold TM, Lim M, Glover PM, Morris PG, Bowtell R, Barnes GR, Brookes MJ (2017) A new generation of magnetoencephalography: room temperature measurements using optically-pumped magnetometers. Neuroimage 149:404–414.

7. Gutteling TP, Bonnefond M, Clausner T, Daligault S, Romain R, Mitryukovskiy S, Fourcault W, Josselin V, Le Prado M, Palacios-Laloy A, Labyt E, Jung J, Schwartz D (2023) A new generation of OPM for high dynamic and large bandwidth MEG: the 4He OPMs—first applications in healthy volunteers. Sensors (Basel) 23:2801.

8. Marhl U et al. (2022) Auditory evoked fields measured simultaneously by OPM- and SQUID-MEG in healthy adults. Neuroimage 247:118827.

9. Tanner Z, Rier L, Fildes J, Reina Rivero G, Schofield H, Marani C, Holmes N, Hill RM, Shah V, Doyle C, Osborne J, Bobela D, Brookes MJ, Boto E (2025) Demonstrating equivalence across magnetoencephalography scanner platforms using neural fingerprinting. Imaging Neurosci 3:IMAG.a.10.

10. Fife AA et al. (1999) Synthetic gradiometer systems for MEG. IEEE Trans Appl Supercond 9:4063–4068.

11. Tierney TM et al. (2024) Adaptive multipole models of optically pumped magnetometer data. Hum Brain Mapp 45:e26596.

12. Mosher JC, Lewis PS, Leahy RM (1992) Multiple dipole modeling and localization from spatio-temporal MEG data. IEEE Trans Biomed Eng 39:541–557.

13. Oostenveld R, Fries P, Maris E, Schoffelen JM (2011) FieldTrip: open source software for advanced analysis of MEG, EEG, and invasive electrophysiological data. Comput Intell Neurosci 2011:156869.

14. Sarvas J (1987) Basic mathematical and electromagnetic concepts of the biomagnetic inverse problem. Phys Med Biol 32:11–22.

15. Nolte G (2003) The magnetic lead field theorem in the quasi-static approximation and its use for magnetoencephalography forward calculation in realistic volume conductors. Phys Med Biol 48:3637–3652.

16. Gallen CC et al. (1994) Intrasubject reliability and validity of somatosensory source localization using a large array biomagnetometer. Electroencephalogr Clin Neurophysiol 90:145–156.

17. Schaefer M et al. (2004) Reproducibility and stability of neuromagnetic source imaging in primary somatosensory cortex. Brain Topogr 17:47–53.

18. Frodl-Bauch T et al. (1997) Dipole localization and test-retest reliability of frequency and duration mismatch negativity generator processes. Brain Topogr 10:3–8.

19. Pang EW et al. (2003) Localization of auditory N1 in children using MEG: source modeling issues. Int J Psychophysiol 51:27–35.

20. Vrba J et al. (1999) Errors in ECD localization with partial sensor coverage. In: Recent advances in biomagnetism: proceedings of the 11th international conference on biomagnetism (Biomag 1998), pp 101–104. Sendai: Tohoku University Press.

21. Paetau R et al. (1995) Auditory evoked magnetic fields to tones and pseudowords in healthy children and adults. J Clin Neurophysiol 12:177–185.

22. Pang EW, Taylor MJ (2000) Tracking the development of the N1 from age 3 to adulthood: an examination of speech and non-speech stimuli. Clin Neurophysiol 111:388–397.

23. Iivanainen J, Stenroos M, Parkkonen L (2017) Measuring MEG closer to the brain: performance of on-scalp sensor arrays. Neuroimage 147:542–553.

24. Zetter R et al. (2018) Requirements for coregistration accuracy in on-scalp MEG. Brain Topogr 31:931–948.

25. Coffey EBJ et al. (2016) Cortical contributions to the auditory frequency-following response revealed by MEG. Nat Commun 7:11070.

